# Visual Complexity, Abstraction, and Human Figuration in Healthcare Wayfinding Symbols: An Eye-Tracking Study

**DOI:** 10.64898/2026.08.08.743673

**Authors:** Azadeh Sharifi Nowghabi, Sara Sharghilavan, Abdolali Bagheri, Morteza Izadifar

## Abstract

Wayfinding in hospitals is often hindered by ineffective signage; however, the cognitive mechanisms of healthcare wayfinding symbols comprehension remain under-researched. This study utilized eye-tracking and spatial gaze mapping to examine how visual complexity, abstraction, and human figuration modulate perception in 40 healthy adults viewing 24 hospital-related healthcare wayfinding symbols. Results indicate that pupil size is a sensitive physiological marker of cognitive load, significantly influenced by visual complexity (*χ*^2^ = 11.32, *p* = .022) and abstraction (*χ*^2^ = 7.49, *p* = .027). Human figuration reduced fixation duration and increased saccade amplitude, facilitating efficient semantic integration. Furthermore, human-centric healthcare wayfinding symbols elicited streamlined gaze trajectories, whereas abstract/complex designs induced chaotic scanpaths. These findings suggest that human figuration acts as a cognitive scaffold, reducing mental effort. We provide evidence-based guidelines for optimizing healthcare wayfinding symbols by prioritizing human body representations and balancing abstraction levels.

**Highlight:**

- Pupil size indexes cognitive load during symbol comprehension.
- Human figuration cuts fixation duration, boosting wayfinding efficiency.
- Abstract symbols increase pupil dilation, raising cognitive load.

## 1. Introduction

Wayfinding in hospitals is persistently challenging due to architectural complexity, frequent spatial changes, inconsistent naming, and poor signage (Tzeng & Huang, 2009; Morag et al., 2016; Suchá et al., 2025). These difficulties disproportionately affect first-time visitors and individuals with physical, cognitive, or visual impairments, leading to confusion, delays, and disrupted access to care (MacKenzie & Krusberg, 1996; Liu et al., 2006; Devlin, 2014; Upadhyay et al., 2022; Kwon et al., 2025). Beyond patient outcomes, such inefficiencies reduce staff productivity and degrade overall healthcare quality (Lee et al., 2020; Jamshidi et al., 2025).

Environmental graphic design, particularly healthcare wayfinding symbols, is critical for intuitive navigation in healthcare settings (Sunyavivat & Boonyachut, 2013; Potter, 2017; Menon et al., 2024). The communicative effectiveness of these symbols depends on key design factors: visual complexity (perceptual density and structural intricacy; Forsythe, 2009), level of abstraction (deviation from realistic depiction, requiring inferential processing; Viola, Chen & Isenberg, 2020; Baker & Kellman, 2018), and the presence of human figuration. Human figuration may activate the observer’s body schema, a dynamic, unconscious representation of the body in space (Gallagher, 2005), thereby facilitating meaning construction through embodied perceptual alignment (Groen et al., 2013). healthcare wayfinding symbols lacking human figuration may lack this scaffolding, potentially increasing reliance on top-down inference and cognitive load.

Therefore, from a cognitive neuroscience perspective, the effectiveness of healthcare wayfinding symbols extends beyond visual appearance and is rooted in the cognitive (including attention and perception) mechanisms engaged during user interaction (Beaufils et al., 2014; Schloss, 2025). Designing effective healthcare wayfinding symbols, therefore, requires a nuanced understanding of how the human brain processes visual information, specifically, how attention is initially directed toward a stimulus and how that stimulus is subsequently perceived and interpreted.

Attention serves as the initial gateway through which information enters the cognitive system (Amso & Scerif, 2015; Wehrle, 2022). In complex and crowded visual environments such as hospitals, a large array of stimuli compete for the user’s cognitive resources, increasing cognitive load (Crosby *et al*., 2001; Beck *et al*., 2010; Rodrigues & Pandeirada, 2019). Attentional mechanisms, both bottom-up, driven by salient visual features such as high contrast, color, or motion, and top-down, guided by the individual’s goals, expectations, and previous knowledge, determine which stimuli are selected for further processing (Katsuki & Constantinidis, 2014; Zheng et al., 2024). Therefore, for a symbol to perform its communicative function of facilitating navigation, it must be both clear and expressive enough to evoke a person’s previous knowledge and cognitively salient enough to focus the person’s attention on it. Hence, Attention can be viewed as not merely a facilitating factor, but a necessary prerequisite for any subsequent understanding (Lindström, 2011; Merritt & Valaris, 2017; Mishra & Prasad, 2025).

Once attention is captured, the process of perception begins (Pashler, 1995; Rensink, 2013; Wilck & Altarriba, 2021). Perception typically involves organizing, interpreting, and assigning meaning to the selected visual information (Pinna & Šķdilters, 2010; Gillebert & Humphreys, 2014; Nisa *et al*., 2023). According to feature integration theory (Treisman & Gelade, 1980), attention acts as a binding mechanism that integrates distinct visual features, such as lines, shapes, and colors, into a coherent whole. In the context of comprehension of healthcare wayfinding symbols, this means that the user must first pay attention to the symbol and then, through organizing and integrating the constituent elements, combine it into a single representation and give it meaning.

Eye tracking technology is a powerful and noninvasive tool for investigating these cognitive processes (Beesley *et al*., 2019). Studies have shown that cognitive processes can be influenced by eye movements. For example, Fixations index attentional focus and encoding (Russo, 2019), with duration reflecting cognitive effort (Irwin, 2013; Liu et al., 2022). Saccades indicate attention shifts (Poole & Ball, 2006; Wolf & Lappe, 2021), gaze trajectory reveals spatiotemporal sampling (Engbert et al., 2015; Zhu et al., 2022), blink rate tracks cognitive load (Walrath et al., 1984; Maffei & Angrilli, 2018), and pupil size indexes mental workload (Alnæs et al., 2014; Oliva, 2019; Gorin et al., 2024).

Despite the established utility of eye tracking in cognitive science, its application in environmental graphic design for evaluating healthcare wayfinding symbols remains limited. A critical gap exists in the lack of multimodal approaches that simultaneously capture overt attentional allocation through eye movements and covert cognitive processing through response time and accuracy measures. Response time serves as a reliable indicator of cognitive load and semantic accessibility (Gwizdka, 2010), and combining it with eye-tracking metrics enables researchers to dissociate whether perception difficulties originate from failures in attentional capture (e.g., delayed first fixation) or from inefficiencies in perceptual and semantic processing (e.g., prolonged fixation duration, pupil dilation). This distinction is essential for developing evidence-based design guidelines that address specific cognitive bottlenecks.

In the present study, we employed a multimodal approach integrating eye tracking and behavioral response time measures to systematically investigate the cognitive processes underlying the perception of healthcare wayfinding symbols. Specifically, we aimed to: (1) characterize the attentional and perceptual mechanisms engaged during symbol interpretation using eye-tracking metrics including fixations, saccades, gaze trajectories, blink rate, and pupil dilation; (2) examine the relationship between these parameters and behavioral outcomes (response time and comprehension accuracy); and (3) identify design features that facilitate or impede efficient attentional capture and semantic interpretation. We hypothesized that healthcare wayfinding symbols with higher visual salience and semantic clarity would elicit shorter time to first fixation, shorter fixation durations, fewer revisits, and faster response times with higher accuracy, whereas ambiguous or complex healthcare wayfinding symbols would produce prolonged fixations, increased revisits, greater pupil dilation, slower response times, and higher error rates. By elucidating the interplay between attention, perception, and comprehension, this study aims to provide empirically grounded insights for developing more intuitive and universally accessible wayfinding systems in healthcare environments.

## 2. Method

### 2.1 Participants

A total of 40 participants (20 females, 20 males; age range: 36-55 years, M = 47.3) were recruited from the healthy general population using convenience sampling. Inclusion criteria were purposively defined to control for potential confounding variables and included: age between 36 and 55 years, right-handedness (to control for hemispheric lateralization), absence of psychiatric or neurological disorders, no history of substance abuse, no hospital-related psychological trauma, and no formal education in visual arts, medical, or paramedical fields. All participants reported normal or corrected-to-normal vision. Based on the visual nature of the task, individuals with significant visual impairments (e.g., color vision deficiency assessed via Ishihara plates, or astigmatism > 1.5 diopters) were excluded. Additionally, participants with a history of neurological conditions affecting oculomotor control or attention (e.g., traumatic brain injury, epilepsy) were excluded to avoid confounding eyetracking measures. Moreover, to ensure high-quality eye-tracking data, standard inclusion thresholds were applied. Participants were included in the final analysis only if they achieved a calibration accuracy of < 0.5° of visual angle during the initial calibration and maintained valid fixation data for at least 75% of the recording duration. Participants who failed to meet these criteria or did not complete the experimental session were excluded. Eventually, all participants provided written informed consent before participation. The study protocol was conducted in accordance with the Declaration of Helsinki and was approved by the Ethics Committee of Isfahan University of the Art (IR.UI.REC.1405.007).

### 2.2 Stimuli

Since this project is part of a larger project, for the current study the stimulus set comprised 24 (out of 60) black and white hospital wayfinding symbols selected to represent common clinical departments, procedures, and diagnostic contexts (Sharifi Noghabi & Bagheri, 2026). Functionally, all symbols served identificational purposes, conveying the identity of clinical services or diagnostic modalities, and were classified into three representational categories: (1) treatment or procedural activities (e.g., injection, infusion), (2) diagnostic tools (e.g., CT scanner, X-ray machine), and (3) anatomical references (e.g., neurology represented via cranial silhouette, cardiology via cardiac silhouette). To control for low-level visual confounds that might influence bottom-up attentional capture, all symbols underwent standardized graphical preprocessing. Color information was removed, and stimuli were rendered in a uniform black (#00000) against a white background (#FFFFFF). Luminance contrast, calculated using the Michelson contrast formula (*Lmax* ― *Lmin*)/(*Lmax* + *Lmin*), was maintained at a constant ratio of 0.85 across all stimuli, as verified using a Konica Minolta CS-200 luminance meter. Symbols were normalized for orientation and spatial alignment, with all stimuli presented at a standardized size of 300 × 300 pixels, subtending approximately 5 ∘ × 5 ∘ of visual angle at a viewing distance of 60 cm. The stimulus set was systematically constructed to vary along two theoretically relevant dimensions: visual complexity and level of abstraction, both of which have been implicated in attentional allocation and semantic processing (Snodgrass & Vanderwart, 1980; Forsythe et al., 2003). Visual complexity was operationalized as the density and arrangement of visual elements, encompassing the number of constituent components, line intricacy, contour density, and perceptual load (Donderi, 2006). Level of abstraction was defined as the degree to which a symbol deviates from a literal, iconic representation of its referent, with abstract symbols requiring greater inferential processing (McDougall et al., 1999). Additionally, the presence versus absence of human body representation was coded as a content-related variable, given its potential to modulate semantic accessibility and attentional engagement (Groen et al., 2013). To establish the validity of these classifications, an independent expert evaluation was conducted. A panel of 50 visual arts specialists (mean years of professional experience = 8.4, *SD* = 3.7) rated each symbol on three dimensions: visual complexity (1 = very simple to 7 = very complex), level of abstraction (1 = highly iconic to 7 = highly abstract), and representational clarity (1 = unclear to 7 = very clear). Inter-rater reliability was assessed using Fleiss’ κ, yielding substantial agreement for complexity (*κ* = 0.78), abstraction (*κ* = 0.74), and clarity (*κ* = 0.81). Several categories of symbols were excluded from the stimulus set to minimize confounding influences. These included safety warning symbols, directional signage (e.g., arrows, stairs), general-purpose informational symbols (e.g., reception), symbols specific to hospital staff or restricted access, and symbols heavily dependent on cultural conventions for interpretation (e.g., pharmacy represented via mortar and pestle). This selection strategy was designed to isolate the core perceptual and cognitive processes underlying symbol comprehension, reducing contextual and cultural bias while maximizing internal validity.

### 2.3 Apparatus

Eye movements were recorded through an SMI-250 device from the Eyelink company and by BeGaze software (version 3.0) and Test Center (version 3.0). The SMI-250 (RED250mobile) is an advanced eye-tracking system that employs video-based infrared technology to record eye movements without direct contact. With a sampling rate of up to 250 Hz, it enables precise measurement of rapid eye movements (saccades) and supports both monocular and binocular tracking. The device offers a spatial accuracy of approximately 0.4° and an RMS spatial resolution of 0.03°, making it suitable for psychological and marketing research. The SMI-250 features multipoint and smart calibration and can be integrated with various experimental software platforms as well as other physiological recording devices, such as EEG, via TTL inputs. Operating at a standard distance of 50–80 cm from the participant, its lightweight and portable design allows use in both laboratory and field settings.

### 2.4 Experimental Procedure

The experiment was composed of two sequential phases, both of which were completed in a single session as you can see the procedure in Fig. 1. This project is part of a larger study. Hence, for this study, we selected 24 hospital symbols for analysis. The hospital symbols were presented in both phases, with the trial order independently randomised for each phase and for each participant to control for order effects and fatigue. Before commencing the experimental phases, participants underwent a standard 9-point calibration procedure for the eye tracker. Following successful calibration, the experimenter initiated the first phase. Throughout both phases, participants were seated approximately 60 cm from the monitor with their head position stabilised using a chin rest to ensure consistent viewing geometry and eyetracking data quality. Phase 1 was designed as a passive free-viewing task intended to capture bottom-up attentional allocation and early perceptual processing uncontaminated by response demands. Each trial began with the presentation of a central fixation cross, which remained on screen for a randomly varying interval between 0.8 and 1.2 seconds. This fixation cross served to reset spatial attention and establish a consistent baseline for pupil diameter measurement. Following the fixation cross, a single symbol was presented at the centre of the screen for a fixed duration of two seconds. During this two-second viewing window, participants were instructed simply to look at the symbol naturally, as if they were encountering it in a hospital environment, and to refrain from making any motor response. After the symbol disappeared, a blank grey screen was shown for 0.5 seconds to minimise visual afterimages before the onset of the next trial. Eye-tracking data, including fixations, saccades, gaze trajectories, pupil diameter, and blink rate, were recorded continuously throughout the entire duration of Phase 1. Phase 2 assessed participants’ previous knowledge of each symbol’s meaning while ensuring that adequate time was available for visual processing before any response was required. Each trial in Phase 2 began with a central fixation cross presented for a random duration between 0.8 and 1.2 seconds, identical to Phase 1. Subsequently, the same symbol used in Phase 1 was displayed at the screen centre for exactly four seconds. Crucially, during this four-second interval, participants were instructed not to respond but rather to attend carefully to the symbol and attempt to retrieve its meaning. This forced viewing period ensured that individual differences in early visual encoding speed did not confound response times. Immediately following the offset of the symbol, a response screen appeared, presenting four options. Participants were instructed to press the correct answer key from the keyboard keys 1 to 4 if they knew the meaning of the symbol and could correctly identify what it represented, and to press the key labeled 5 if they did not know the meaning of the symbol. No time limit was imposed on this response, but participants were encouraged to answer as accurately as possible. Following the keypress, a blank screen was presented for 0.5 seconds before the next trial commenced. As in Phase 1, eye-tracking data were recorded continuously during the four-second symbol viewing window in Phase 2, capturing realtime attentional and cognitive processes during semantic retrieval. After the response, eye tracking continued to record but no critical data were analysed during the blank interval. The entire experimental session concluded automatically after the last trial of Phase 2, at which point all behavioural and eye-tracking data were saved for offline analysis.

**Fig. 1.**
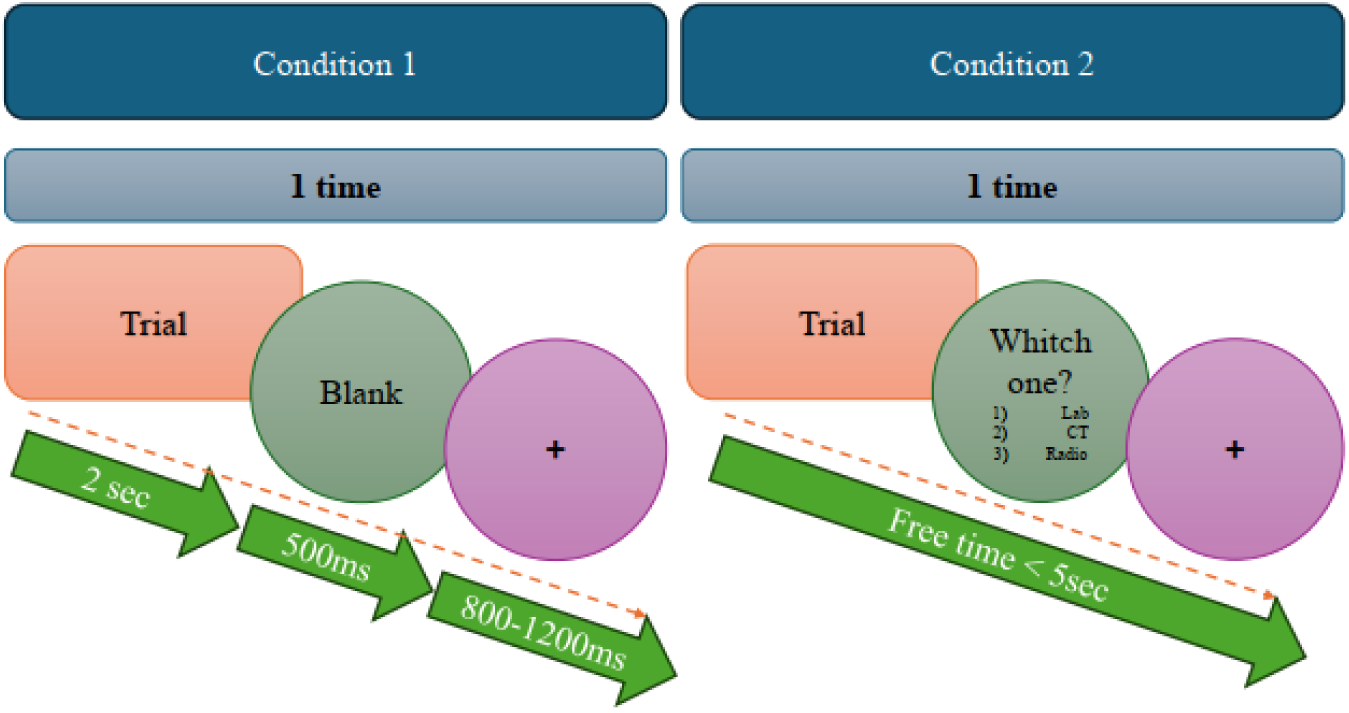
Experimental Procedure

### 2.5 Data analysis

Data preprocessing and outlier removal were performed using Python (Version 3.10) with the pandas (Version 2.0.3) and numpy (Version 1.24.3) libraries, while the pathlib module was used for cross-platform file path management. Raw eye-tracking data, initially exported as Excel files, were imported into the analysis environment, and trials with excessive data loss exceeding 25% of the total stimulus duration were excluded. Outliers were identified and removed using the interquartile range (IQR) method, defined as values falling below Q1 − 1.5 × IQR or above Q3 + 1.5 × IQR for each eye-tracking metric, including fixation duration, saccade amplitude, and pupil size.

To identify spatial clusters of fixations reflecting concentrated visual attention, the Density-Based Spatial Clustering of Applications with Noise (DBSCAN) algorithm was implemented using the sklearn.cluster module (Version 1.3.0). DBSCAN was selected for its ability to identify arbitrarily shaped clusters without requiring the number of clusters to be specified a priori, as well as its robustness to outliers (Ester et al., 1996). Fixation coordinates (x, y pixels) were extracted for each trial and each participant, and the DBSCAN algorithm was applied with an epsilon (ε) of 50 pixels (corresponding to approximately 1.5° of visual angle at the 60 cm viewing distance) and a minimum sample size (min_samples) of 5, both determined empirically through a grid search. Points not assigned to any cluster were classified as noise, representing sporadic fixations not reflecting sustained attention. For each identified cluster, the cluster centroid coordinates weighted by fixation duration, the number of fixations within the cluster, and the total dwell time were computed.

Following cluster analysis, continuous density maps of visual attention (gaze heatmaps) were generated using Kernel Density Estimation (KDE) via the scipy. stats. gaussian_kde function (SciPy Version 1.11.1). KDE provides a smooth, non-parametric estimate of the probability density function of fixation locations (Caldara & Miellet, 2011). A Gaussian kernel was applied with a bandwidth determined by Scott’s rule, calculated as bandwidth = n^(−1/(d+4)), where n represents the number of fixation points and d represents the dimensionality (d = 2 for spatial coordinates). Fixation durations were used as weights in the density estimation, ensuring that longer fixations contributed more to the density surface than brief, transient fixations. Heatmaps were generated for each symbol individually and aggregated across all stimuli to reveal general viewing patterns. All visualizations, including fixation cluster plots (with cluster centroids marked as circles using matplotlib.patches.Circle) and gaze heatmaps, were generated using matplotlib. pyplot (version 3.7.2), with warnings suppressed using the warnings module to maintain output clarity. All visualizations were exported as high-resolution (300 dpi) PNG and PDF files for subsequent reporting.

Finally, statistical comparisons were conducted to examine differences in attentional allocation between symbols differing in design features, such as visual complexity and presence versus absence of human body representation. For each symbol, the number of fixation clusters (indexing attentional dispersion), mean cluster dwell time (indexing sustained attention), and spatial entropy of clusters (indexing attentional uncertainty, calculated using Shannon’s entropy across cluster centroids) were extracted.

## 3. Results

In this section, descriptive statistics for all eye-tracking parameters, stratified by visual complexity, abstraction, and human figuration, are first presented. To examine the effects of these symbol features on oculomotor behavior, a series of Kruskal-Wallis tests were then performed. Subsequently, to determine the unique contribution of each visual property to key cognitive indicators, multiple linear regression analyses were conducted on pupil size, fixation duration, and saccade amplitude.

Table 1 shows descriptive statistics for age, sex, handedness, vision status, exclusion criteria, and eye-tracking data quality. All participants were right-handed, had normal or corrected-to-normal vision, and met stringent eye-tracking quality thresholds (calibration accuracy < 0.5° of visual angle, valid fixation data > 75% of recording duration). None of the participants met any of the predefined exclusion criteria (visual arts education, medical/paramedical education, neurological/psychiatric disorders, color vision deficiency, or astigmatism > 1.5 diopters). Values in parentheses represent percentages of the total sample where applicable.

**Table 1.** Demographic, clinical, and eye-tracking quality characteristics.

| Characteristic | Category / Statistic | Value |
| --- | --- | --- |
| Participants (N) | Total | 40 |
|  | Female | 20 (50%) |
|  | Male | 20 (50%) |
| Age (years) | Range | 36-55 |
|  | Mean (M) | 47.3 |
|  | Standard Deviation (SD) | - |
| Handedness | Right-handed | 40 (100%) |
| Vision | Normal or corrected-to-normal | 40 (100%) |
| Exclusion criteria applied | Visual arts education | 0 |
|  | Medical/paramedical education | 0 |
|  | Neurological/psychiatric disorders | 0 |
|  | Color vision deficiency | 0 |
|  | Astigmatism > 1.5 D | 0 |
| Eye-tracking data quality | Calibration accuracy < 0.5° | 40 (100%) |
|  | Valid fixation data > 75% | 40 (100%) |

Table 2 presents the means, standard deviations (SD), and standard errors (SE) for six eye-tracking metrics, blink count, blink rate, fixation count, fixation duration (ms), saccade count, and pupil size (arbitrary units), stratified by three symbol properties: visual complexity (five levels: very low, low, moderate, high, very high), abstraction (four levels: very low, low, moderate, very high), and human figuration (two levels: yes, no). For each combination of eye-tracking metric and symbol property, the mean and associated standard error are reported in the Statistic and Std. Error columns, respectively, with the standard deviation provided directly below each mean.

**Table 2.**
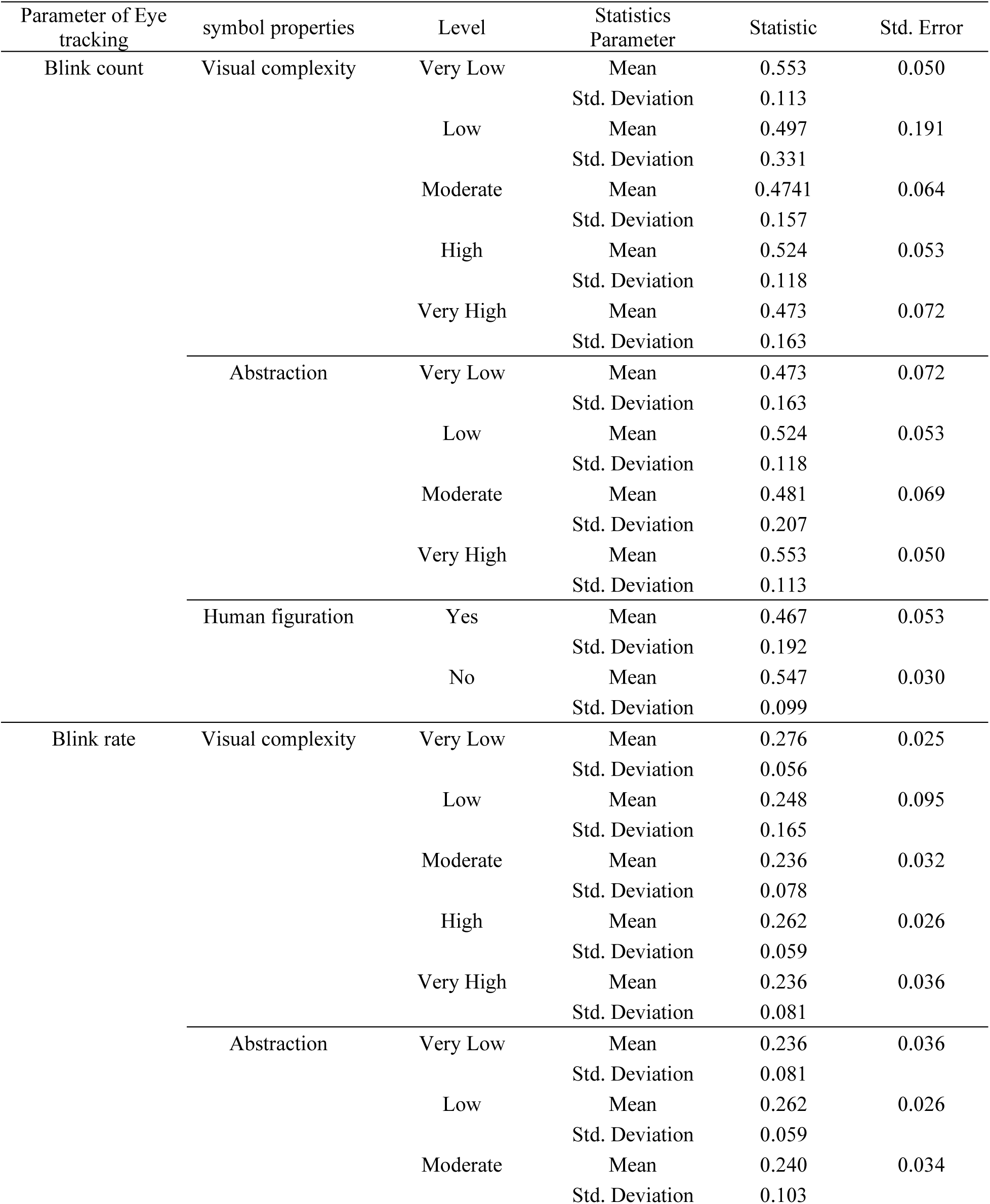
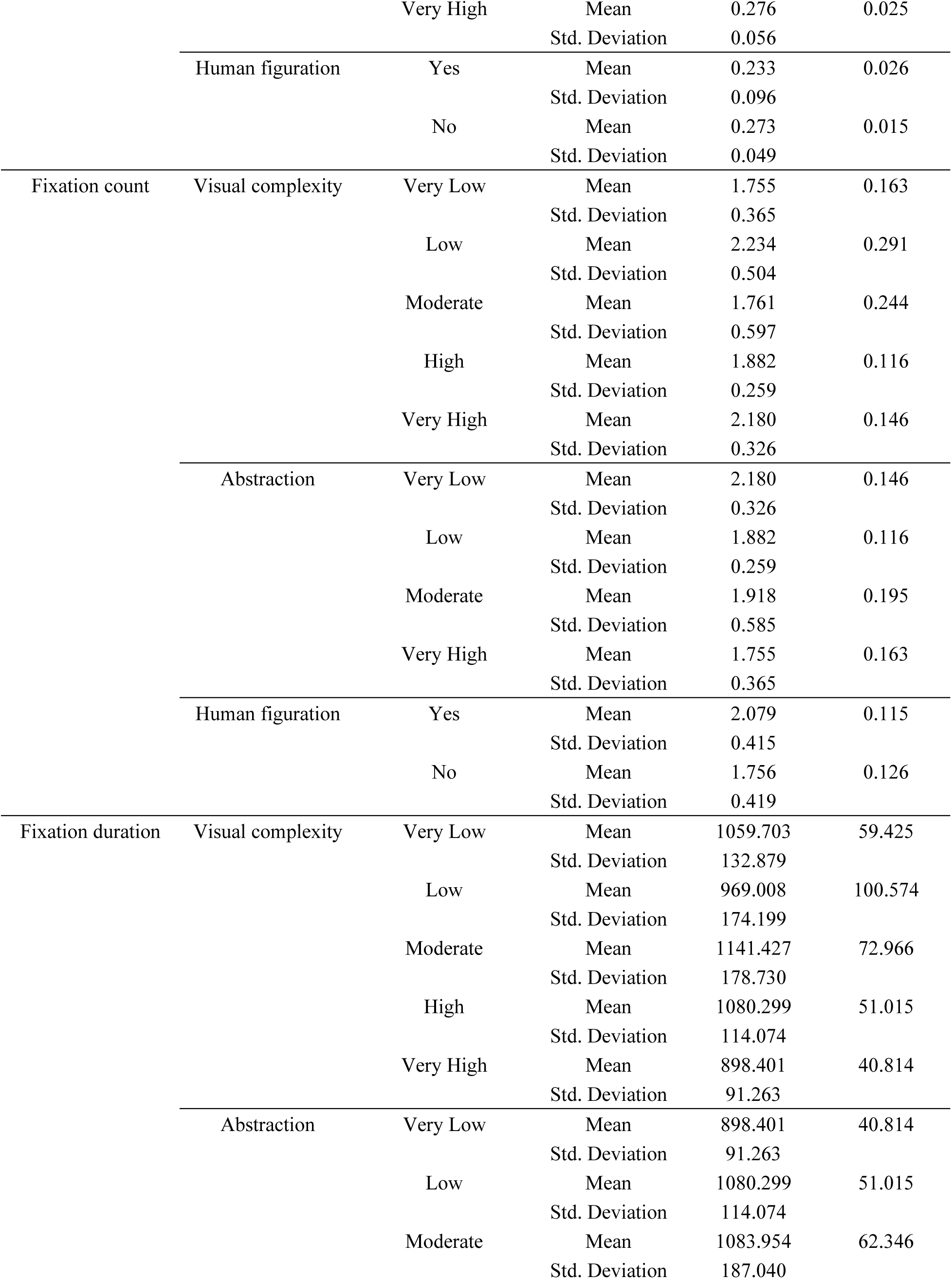
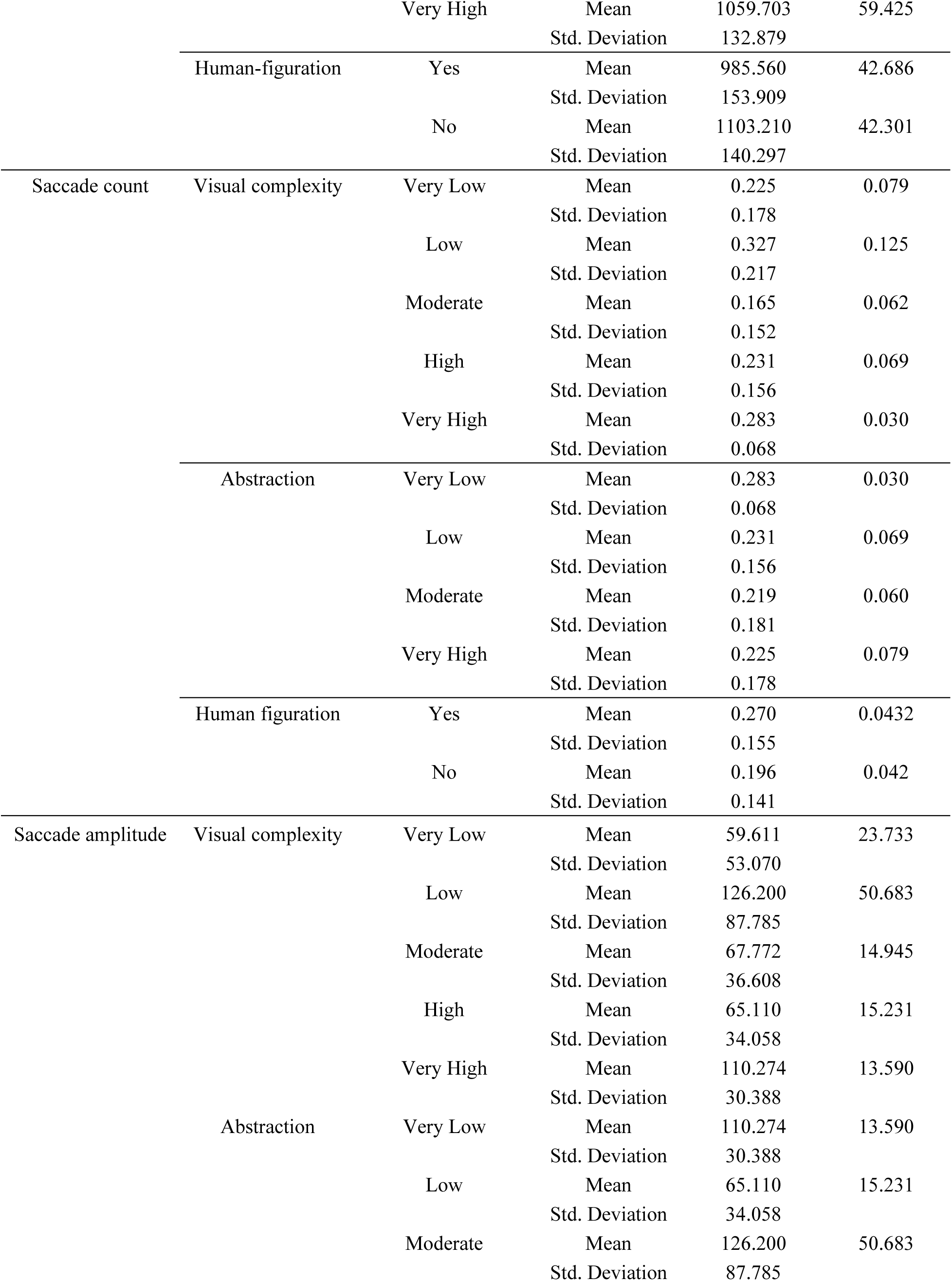
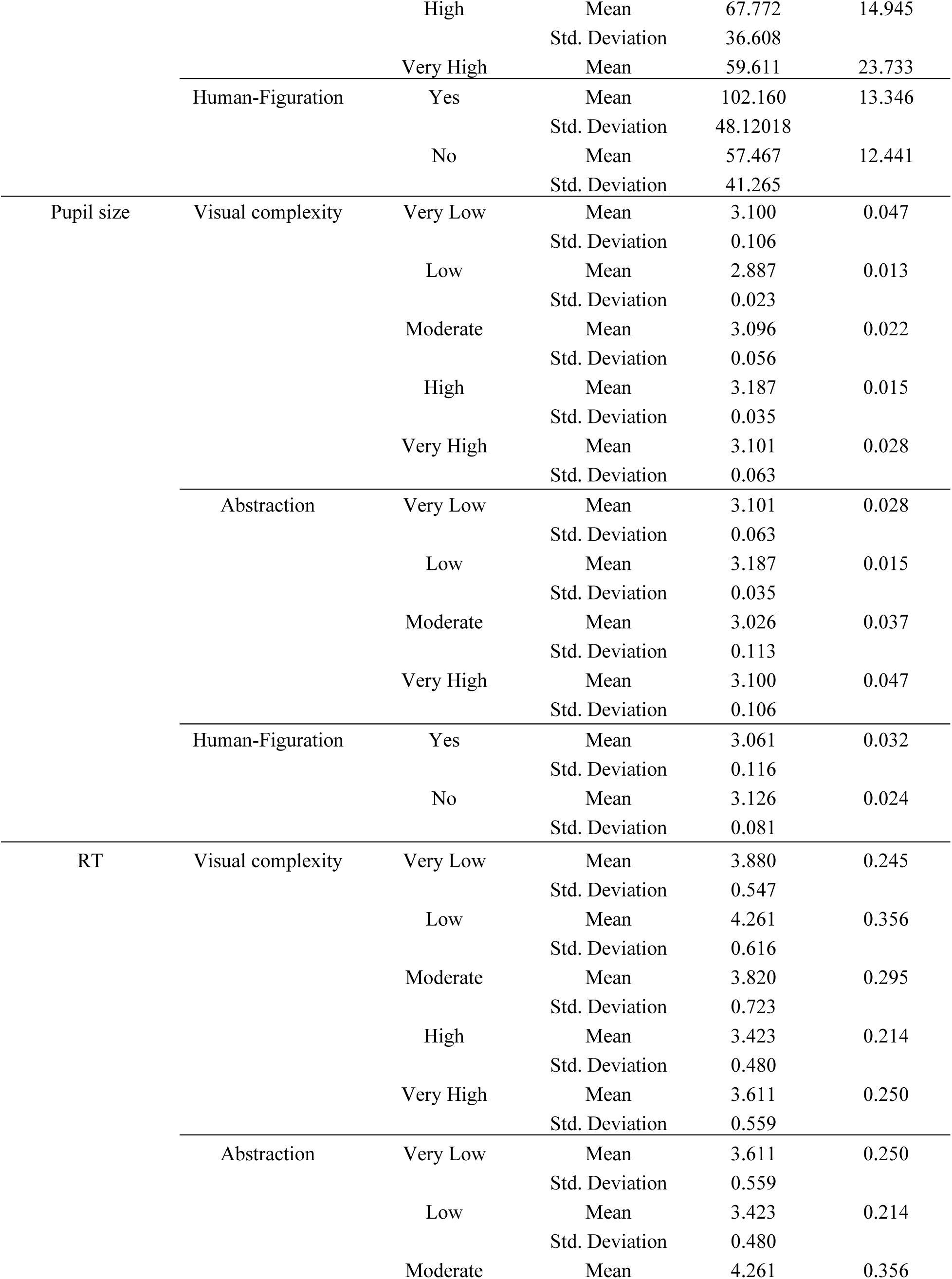
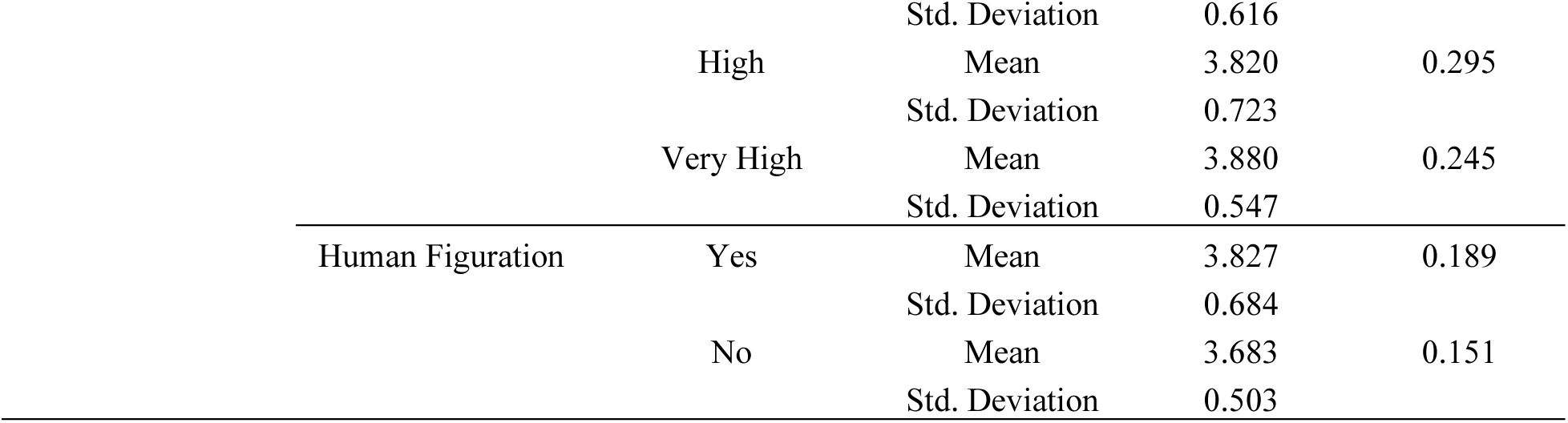
Descriptive statistics of eye-tracking parameters across visual complexity, abstraction, and human figuration.

In fig.2, for each symbol, the left panel presents the original stimulus, the middle panel displays the gaze heatmap generated using Kernel Density Estimation (KDE), and the right panel illustrates fixation trajectories and spatial fixation clusters identified through the DBSCAN algorithm. Warmer colors in the heatmaps indicate regions with higher fixation density and longer dwell time. Overall, participants exhibited concentrated visual attention toward semantically salient central elements of the symbols, while visually complex or anatomically detailed symbols elicited more dispersed fixation patterns and longer scan paths. Symbols characterized by lower visual complexity and clearer structural organization demonstrated more centralized fixation clusters and reduced attentional dispersion, suggesting more efficient perceptual processing and semantic accessibility.

**Fig 2:**
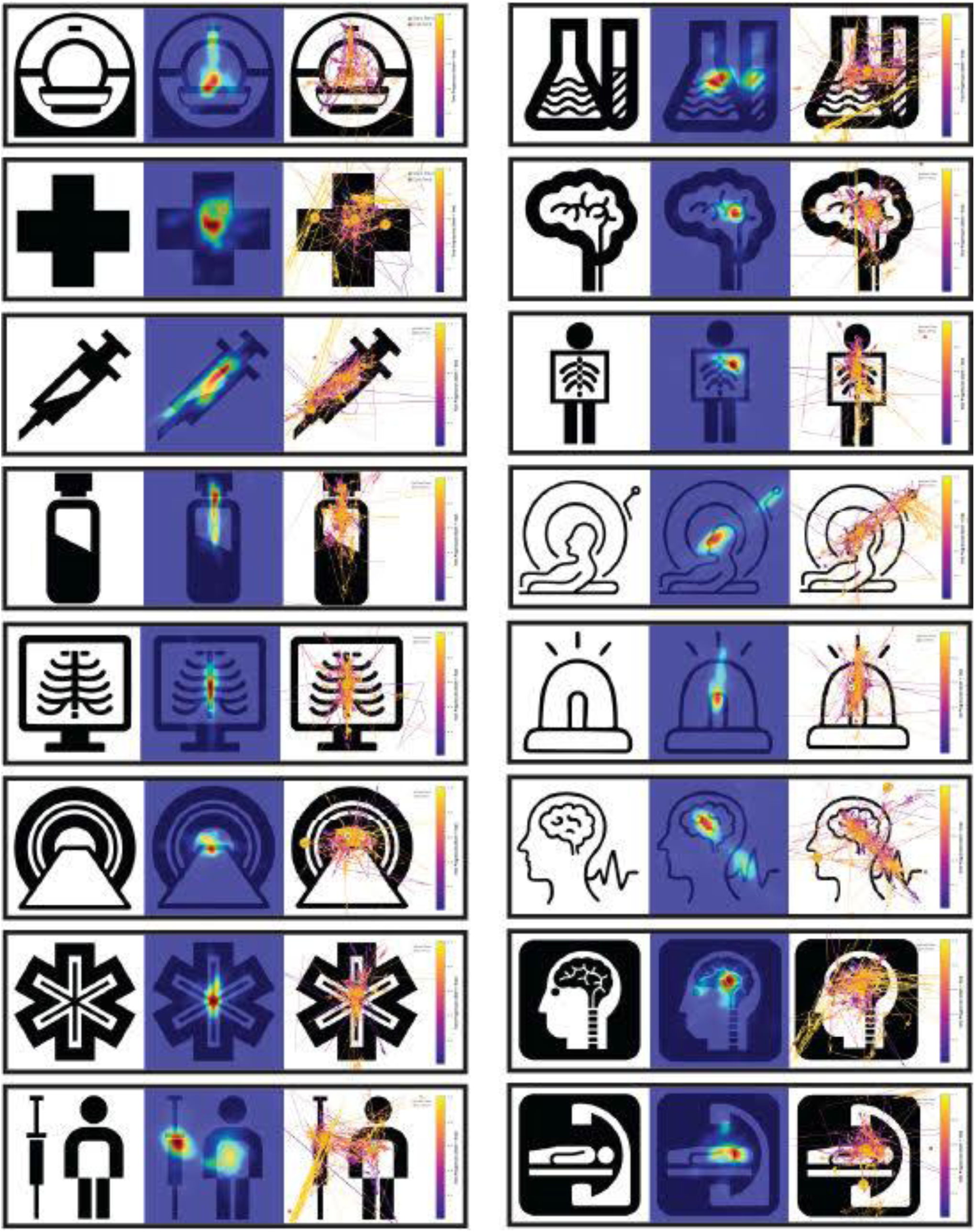
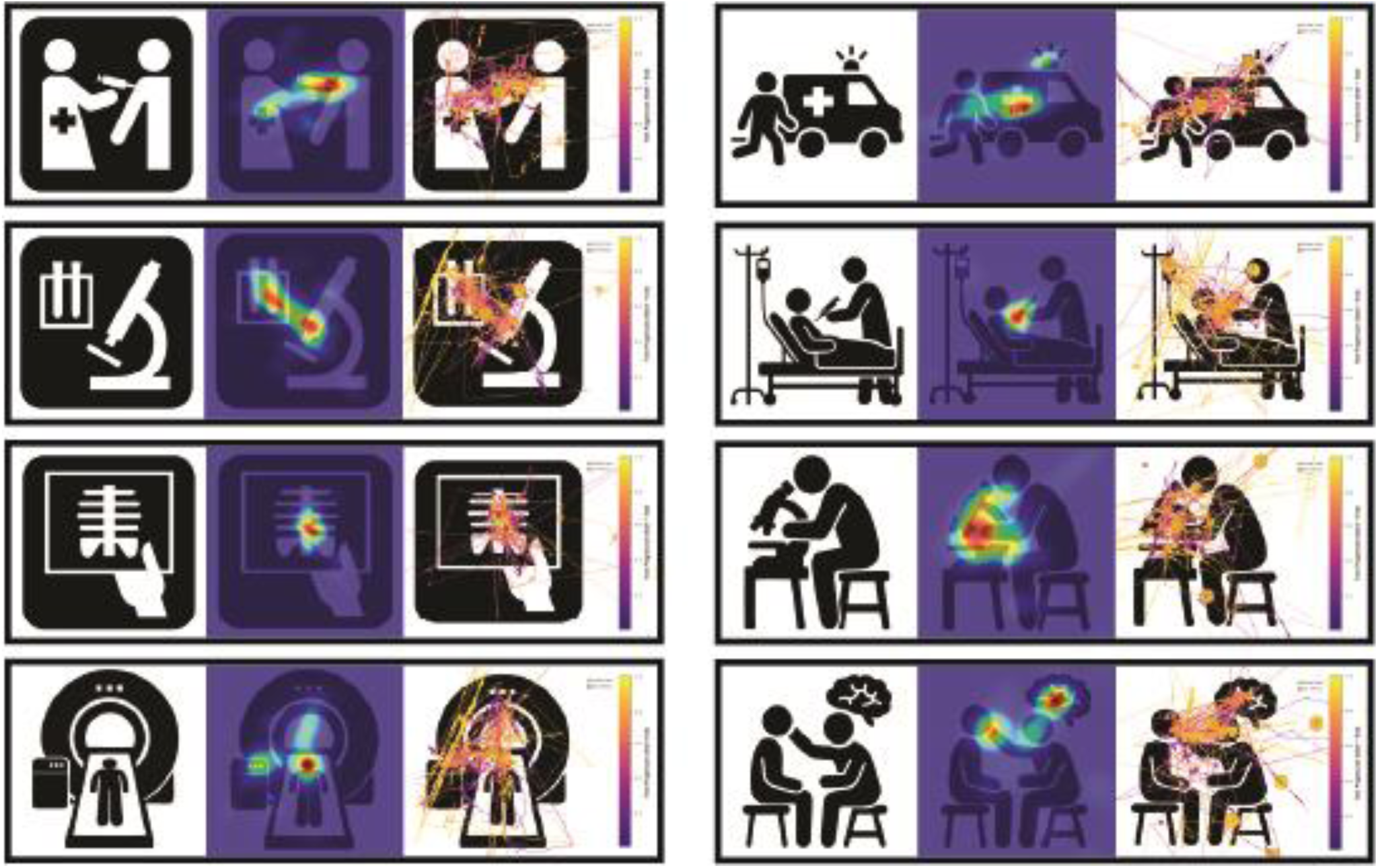
Symbols, heat map, and gaze trajectory of each symbol

As shown in Table 3, chi-square analyses revealed that visual complexity and abstraction significantly affected pupil size (χ² = 11.32, df = 4, p = .022; χ² = 7.49, df = 4, p = .027, respectively), whereas other metrics were not significant. Average fixation duration approached significance for both visual complexity and abstraction (p= .069). Human figuration significantly influenced average fixation duration (χ² = 4.21, df = 1, p = .040) and saccade amplitude (χ² = 4.96, df = 1, p = .026), with saccade count nearing significance (p= .077). Blink count, blink rate, fixation count, and pupil size were unaffected by human figuration. These findings suggest that pupil size is particularly responsive to visual complexity and abstraction, whereas human figuration modulates gaze dynamics, including fixation duration and saccade amplitude.

**Table 3.** Results of the Kruskal-Wallis tests comparing eye-movement characteristics across symbol properties.

| symbol properties | Dependent Variable | $\chi^2$ | df | $p$ | Significance |
| --- | --- | --- | --- | --- | --- |
| Visual complexity | Blink count | 1.91 | 4 | 0.752 | N.S. |
|  | Blink rate | 1.85 | 4 | 0.757 | N.S. |
|  | Fixation count | 5.42 | 4 | 0.252 | N.S. |
|  | Average fixation duration | 8.78 | 4 | 0.069 | Near significant |
|  | Saccade amplitude | 5.98 | 4 | 0.200 | N.S. |
|  | Saccade count | 4.92 | 4 | 0.295 | N.S. |
|  | Pupil size | 11.32 | 4 | 0.022 | Significant |
|  | RT | 3.37 | 4 | 0.498 | N.S. |
| Abstraction | Blink count | 1.80 | 4 | 0.774 | N.S. |
|  | Blink rate | 1.76 | 4 | 0.767 | N.S. |
|  | Fixation count | 5.08 | 4 | 0.281 | N.S. |
|  | Average fixation duration | 7.65 | 4 | 0.105 | N.S. |
|  | Saccade amplitude | 5.31 | 4 | 0.260 | N.S. |
|  | Saccade count | 1.89 | 4 | 0.747 | N.S. |
|  | Pupil size | 7.49 | 4 | 0.027 | Significant |
|  | RT | 3.28 | 4 | 0.512 | N.S. |
| Human figuration | Blink count | 1.86 | 1 | 0.173 | N.S. |
|  | Blink rate | 2.01 | 1 | 0.156 | N.S. |
|  | Fixation count | 2.72 | 1 | 0.099 | N.S. |
|  | Average fixation duration | 4.21 | 1 | 0.040 | Significant |
|  | Saccade amplitude | 4.96 | 1 | 0.026 | Significant |
|  | Saccade count | 3.13 | 1 | 0.077 | Near significant |
|  | Pupil size | 2.18 | 1 | 0.140 | N.S. |
|  | RT | 3.08 | 1 | 0.079 | N.S. |

Before multiple linear regression, assumptions were tested. Normality of residuals was assessed using the Shapiro-Wilk test (all p > .05). Homoscedasticity was confirmed via Breusch-Pagan test (all p > .05). Multicollinearity was examined using Variance Inflation Factor (VIF), with all values below 2.5, indicating no significant multicollinearity among predictors This allowed us to proceed with multiple linear regression analysis.

As shown in Table 4, a multiple linear regression analysis was conducted to examine whether visual complexity, abstraction, and human figuration predict pupil size. The overall model was statistically significant, *F*(3, 36) = 5.898, *p* = .005, explaining approximately 47% of the variance in pupil size (*R²* = .47).

**Table 4.** Results of Multiple Linear Regression Analysis Predicting Pupil Size.

| Predictor | B | SE | $\beta$ | t | p | Model fit |
| --- | --- | --- | --- | --- | --- | --- |
| Constant | 2.528 | 0.186 | - | 13.560 | < .001 |  |
| Visual Complexity | 0.079 | 0.032 | 1.005 | 2.435 | .024* |  |
| Abstraction | 0.035 | 0.036 | 0.461 | 0.986 | .336 |  |
| Human Figuration | 0.135 | 0.050 | 0.605 | 2.693 | .014* |  |
| | | | | | | $R^2 = .47$ |
| | | | | | | $F(3, 36) = 5.898$ |
| | | | | | | $p = .005$ |

*Dependent variable: pupil size.*
*B = unstandardized coefficient; SE = standard error; $\beta$ = standardized coefficient.*
*$R^2$ = coefficient of determination.*
*Significant predictors are marked with \* ( $p < .05$ ).*

Among the predictors, visual complexity (*B* = 0.079, *p* = .024) and human figuration (*B* = 0.135, *p* = .014) were significant positive predictors of pupil size, whereas abstraction (*p* = .336) was not statistically significant.

As shown in Table 5, a multiple linear regression analysis was conducted to examine whether visual complexity, abstraction, and human figuration predict fixation duration. The overall model was not statistically significant, *F*(3,36) = 1.906, *p* = .161, explaining approximately 22.2% of the variance in fixation duration (*R*^2^ = .222).

**Table 5.** Results of Multiple Linear Regression Analysis Predicting Fixation Duration.

| Predictor | B | SE | $\beta$ | t | p | Model Fit |
| --- | --- | --- | --- | --- | --- | --- |
| Constant | 550.573 | 330.061 | - | 1.668 | .111 |  |
| Visual Complexity | 51.163 | 57.480 | 0.445 | 0.890 | .384 |  |
| Abstraction | 71.040 | 63.120 | 0.638 | 1.125 | .274 |  |
| Human Figuration | 73.732 | 88.611 | 0.226 | 0.832 | .415 |  |
| Model Summary |  |  |  |  |  | R<sup>2</sup> = .222 |
|  |  |  |  |  |  | F(3, 36) = 1.906 |
|  |  |  |  |  |  | p = .161 |

*Dependent variable: fixation duration*
*B = unstandardized coefficient; SE = standard error; $\beta$ = standardized coefficient.*
*$R^2$ = coefficient of determination.*
*Significant predictors are marked with \* ( $p < .05$ ).*

Among the predictors, none reached statistical significance: visual complexity (*B* = 51.163, *p* = .384), abstraction (*B* = 71.040, *p* = .274), and human figuration (*B* = 73.732, *p* = .415). The constant term was also not significant (*B* = 550.573, *p* = .111).

These findings suggest that fixation duration is not significantly predicted by low-level image attributes such as visual complexity, abstraction, or human figuration in the current sample.

As shown in Table 6, a multiple linear regression analysis was conducted to examine whether visual complexity, abstraction, and human figuration predict saccadic amplitude. The overall model was not statistically significant, *F*(3, 36) = 2.031, p = .142, explaining approximately 23.4% of the variance in saccadic amplitude (*R²* = .234). Among the predictors, none reached statistical significance: visual complexity (*B* = −13.417, p = .441), abstraction (*B* = −12.746, p = .505), and human figuration (*B* = −40.082, p = .144). The constant term was significant (*B* = 220.268, p= .036), indicating a baseline saccadic amplitude when all predictors are zero.

**Table 6.** Results of Multiple Linear Regression Analysis Predicting Saccade Amplitude.

| Predictor | B | SE | $\beta$ | t | p | Model Fit |
| --- | --- | --- | --- | --- | --- | --- |
| Constant | 220.268 | 98.103 | - | 2.245 | .036 |  |
| Visual Complexity | -13.417 | 17.085 | -0.390 | -0.785 | .441 |  |
| Abstraction | -12.746 | 18.761 | -0.382 | -0.679 | .505 |  |
| Human Figuration | -40.082 | 26.338 | -0.411 | -1.522 | .144 |  |
| Model Summary | | | | | | $R^2 = .234$ |
| | | | | | | $F(3, 36) = 2.031$ |
| | | | | | | $p = .142$ |

*Dependent variable: saccade amplitude.*
*B = unstandardized coefficient; SE = standard error; $\beta$ = standardized coefficient.*
*$R^2$ = coefficient of determination.*
Significant predictors are marked with \* ( $p < .05$ ).

## 4. Discussion

The present study employed a multimodal approach integrating eye-tracking metrics with spatial gaze mapping to investigate how visual complexity, abstraction, and human figuration modulate attention and perception in hospital symbol comprehension. Our findings offer preliminary evidence for the distinction between attentional capture and perceptual integration as two sequential but interdependent cognitive processes.

According to the visual saliency (Ullah et al., 2020) and feature integration (Quinlan, 2003) theories, participants tended to gaze predominantly at semantically informative regions, which may suggest that attentional selection can be guided by both low-level features and cognitive relevance (Henderson, Malcolm & Schandl, 2009). Simpler symbols tended to produce centralized fixations, shorter gaze trajectories, and lower spatial entropy, which may reflect reduced cognitive load. Complex or anatomically detailed symbols generated distributed fixations and increased revisits, indicating higher perceptual uncertainty, consistent with prior research (Ruddle & Lessels, 2006; Bauer & Ludwig, 2019). Human forms facilitated attentional organization (Rousek & Hallbeck, 2011), potentially activating embodied representations and body-schema mechanisms. Heatmaps revealed a central viewing bias, confirming attentional capture across all conditions. However, post-capture distribution differed by category: treatment symbols showed a bimodal distribution around body parts and tools; diagnostic symbols focused on mechanically distinctive components; anatomical symbols with human figuration elicited broader exploration than isolated organs, suggesting that the human body frame provides an attentional scaffold.

Based on feature integration theory (Treisman & Gelade, 1980) and the literature on bottom-up versus top-down attention (Katsuki & Constantinidis, 2014), these design properties exerted differential effects on attentional capture versus semantic interpretation. Pupil size emerged as a potentially sensitive physiological marker of cognitive load, significantly affected by both visual complexity and abstraction. Regression analysis indicated that visual complexity and human figuration positively predicted pupil dilation, whereas abstraction alone was non-significant in the regression model, despite showing a significant effect in the Kruskal–Wallis test. This pattern suggests that abstraction shares variance with visual complexity, and its influence diminishes when both predictors are analyzed simultaneously. Furtheremore, this finding is broadly consistent with literature linking pupil dilation to mental effort, suggesting that complexity and abstraction may primarily impose load (Alnæs et al., 2014; Oliva, 2019).

Human figuration appeared to be particularly influential, significantly reducing fixation duration and increasing saccade amplitude, indicating more efficient visual scanning and faster semantic integration. Gaze trajectory analysis revealed that human-figuration symbols tended to produce smooth, linear scanpaths with few revisits, whereas abstract symbols without human figures produced chaotic, looping trajectories with multiple regressions. Importantly, although human figuration reduced fixation duration and increased saccade amplitude, regression analysis revealed that it positively predicted pupil dilation. This suggests that human bodies may function as a cognitive scaffold: they reduce the need for repeated fixations and support efficient semantic integration (as indicated by shorter fixation durations), yet simultaneously elicit heightened early attentional engagement (reflected in increased pupil dilation), likely due to their biological and perceptual salience (Groen et al., 2013).

In contrast to hypotheses, blink rate, blink count, and fixation count did not differ significantly across conditions. This suggests that cognitive load may not have reached the threshold required to modulate spontaneous eyeblink rate (Maffei & Angrilli, 2018), or that the fixed viewing durations were insufficient. The dissociation between fixation count (unchanged) and fixation duration (varying with human figuration) is methodologically important, supporting the use of multimodal metrics that capture not merely where and how often people look, but how hard the visual system is working.

Descriptive statistics revealed an inverted-U relationship between visual complexity and fixation duration, with the longest fixations for moderate complexity and shorter fixations for both very low and very high complexity. This non-linear pattern suggests that very low complexity allows rapid processing, moderate complexity is most effortful, and very high complexity may induce satisficing, reducing fixation duration but likely also reducing comprehension accuracy. Response time was not significantly affected by any design property, possibly reflecting the forced viewing period that ensured adequate encoding time, meaning that response time indexed post-perceptual decision-making rather than early visual processing speed.

These findings translate into actionable design guidelines for evidence-based environmental graphic design in healthcare settings. Hospital symbols should incorporate human body representations wherever semantically appropriate, as human figuration reduces fixation duration, increases saccade amplitude, and produces efficient linear gaze trajectories. Gratuitous visual complexity should be minimized, as high complexity increases cognitive load without improving communicative clarity and produces fragmented scanpaths. Moderate levels of abstraction are optimal because overly abstract symbols increase inferential load and attentional uncertainty. Gaze trajectory analysis should be used diagnostically in the design process: linear, non-revisiting scanpaths indicate cognitive efficiency, whereas chaotic, looping trajectories indicate a need for redesign. Finally, the inverted-U relationship between complexity and fixation duration suggests that designers should aim for moderate complexity that provides sufficient discriminative features without exceeding working memory capacity.

## 5. Conclusion

In conclusion, this study suggests that pupil size may be a particularly sensitive eye-tracking metric for detecting cognitive load differences during hospital symbol comprehension, with visual complexity and the presence versus absence of human body representation may modulate mental effort during perceptual integration. Fixation duration may provide complementary information about semantic integration difficulty, while heatmaps and gaze trajectories reveal qualitatively distinct attentional strategies: human figuration produces smooth, linear scanpaths with few revisits, whereas abstract or highly complex symbols produce chaotic, looping trajectories with multiple regressions. Human figuration facilitates faster, more efficient processing by reducing fixation duration and increasing saccade amplitude, despite unexpectedly increasing pupil dilation, suggesting that cognitive benefits emerge during semantic integration rather than early perceptual encoding These findings offer preliminary, empirically grounded guidelines that could be further validated for evidence-based environmental graphic design: hospital symbols should incorporate human body representations where semantically appropriate, minimize gratuitous visual complexity, maintain moderate levels of abstraction, and be validated using gaze trajectory analysis before deployment. By bridging cognitive science and environmental graphic design, this multimodal approach offers a potential framework for developing intuitive, accessible, and universally usable wayfinding systems in healthcare environments, ultimately reducing patient confusion, improving staff efficiency, and enhancing the overall quality of care.

### Limitation and Future Direction

Regarding limitations, the present study used only grayscale symbols and did not include colored symbols. This may limit the generalizability of the findings, as both grayscale and colored symbols are used in hospital wayfinding systems. In addition, the sample was restricted to healthy adults and did not include older adults or individuals with cognitive impairments, who represent important users of hospital navigation systems. Future research should therefore examine colored symbols and extend the investigation to these populations to provide more comprehensive evidence for the ergonomic design of healthcare signage.

## Acknowledgments

The authors gratefully acknowledge the valuable collaboration of participants.

## Consent to publish

Written informed consent for publication was obtained from all participants or, where applicable, from their parents or legal guardians.

## Consent to Participate

Written informed consent was obtained from all individual participants.

## AI Declaration

During the preparation of this work, the authors used Chat GPT for improving language and readability. After using this tool, the authors reviewed and edited the content as needed and take full responsibility for the content of the published article.

## Authors’ Contributions

**Azadeh Sharifi Nowghabi:** Conceptualization, Methodology, Investigation, Data curation, Visualization, Resources, Software & Writing-Review & Editing.

**Sara Sharghilavan:** Supervision, Conceptualization, Methodology, Investigation, Formal Analysis, Writing-Original Draft, and Writing-Review & Editing.

**Abdolali Bagheri:** Project Administration, Conceptualization, Investigation, Validation, Visualization.

**Morteza Izadifar:** Supervision, Conceptualization, Methodology, Validation, Formal Analysis, and Writing-Review & Editing.

